# A new insight into MYC action: control of RNA polymerase II methylation and transcription termination

**DOI:** 10.1101/2022.02.17.480813

**Authors:** Fiorella Scagnoli, Alessandro Palma, Annarita Favia, Claudio Scuoppo, Barbara Illi, Sergio Nasi

## Abstract

A common catastrophic event in most human cancers is deregulation of MYC, a multifunctional transcription factor that controls gene expression in partnership with MAX and drives key biological mechanisms of the cell. Restraining its activity impairs cancer cell features and prevents tumor development, as shown by Omomyc - a 90 amino acid mini-protein interfering with MYC activity. MYC regulates many aspects of transcription by RNA polymerase II (RNAPII), such as activation, pause release, and elongation. That it may have a role in transcription termination as well is suggested by our finding of an interaction between MYC and the Protein Arginine Methyltransferase 5 (PRMT5), which catalyzes symmetrical dimethylation of RNAPII at the arginine residue R1810 (R1810me2s) allowing proper termination and splicing of transcripts. Here we show that MYC overexpression strongly increases R1810me2s, while the concomitant expression of Omomyc or a MYC-specific shRNA counteracts this capacity. Omomyc impairs as well Serine 2 phosphorylation in the RNAPII carboxyterminal domain, a modification that sustains transcript elongation and is enhanced by MYC. By displacing MYC on DNA, Omomyc reshapes RNAPII distribution along genes, leading to greater occupancy of promoter and termination sites. It is unclear how this may affect expression of the variety of genes that control metabolic, biosynthetic, and other pathways and are up or down regulated upon MYC inhibition. Genes belonging to a signature of direct MYC targets are instead strongly downregulated following MYC inhibition, with a weak correlation with RNAPII occupancy at promoters. Our data point to a MYC/ PRMT5/RNAPII axis that controls termination via RNAPII dimethylation (R1810me2s) and may contribute to fine-tune the expression of genes altered by MYC overexpression in cancer cells. It remains to be seen which role this may have in tumor development and maintenance.

## Introduction

Deregulated expression of the transcription factor MYC is a crucial event in a wide variety of cancer cells. Deregulation is usually due to overexpression and affects many transcriptional networks controlling key cell functions like proliferation, metabolism, stemness maintenance, pre-mRNA splicing. For instance, MYC maintains splicing fidelity during lymphomagenesis by upregulating transcription of the core small nuclear ribonucleoprotein particle assembly genes, including PRMT5, which methylates Sm proteins (Koh et al. 2015). Through its bHLHZip domain, MYC dimerizes with MAX and binds to promoter proximal regions of genes. It is unclear whether it mainly acts as a universal amplifier of the transcriptional program running within a given cell, or as a specific activator of a distinct set of target genes (Guccione et al. 2006; Lin et al. 2012; Nie et al. 2012; Perna et al. 2012; Sabò et al. 2014; Walz et al. 2014; Wolf et al. 2015). MYC inhibition may represent an effective strategy against a variety of cancers, as clearly shown by studies with Omomyc, a MYC derived, 90 amino acid mini-protein that affects MYC binding to promoters and restrains MYC function in cancer cells and *in vivo* models (Soucek et al. 1998; Soucek et al. 2008, Savino et al. 2011; Galardi et al, 2016; Beaulieu et al. 2019; Daffy et al. 2021). MYC enhances transcriptional pause release and elongation upon binding to proteins like CDK9 (the kinase subunit of p-TEFb) and SPT5, which represent some of the proteins and protein complexes that interact with the RNAPII carboxyl terminal domain (CTD) and control transcription, mRNA processing, and nucleosome modifications (Rahl et al. 2010; Rahl et. al. 2014; Poole and van Riggelen 2017; Kress et al. 2015; Lorenzin et al. 2016; Baluapuri et al. 2019). The CTD contains 52 tandem repeats of the consensus sequence N-Tyr1-Ser2-Pro3-Thr4-Ser5-Pro6-Ser7-C, and several Arg (R) and Lys (K) residues that are targets of post-translational modifications - phosphorylation, ubiquitination, methylation - which regulate distinct steps of transcription (Phatnani et al. 2006; Eick et al. 2013; Napolitano et al. 2014; Zhao et al. 2016). The TFIIH complex and p-TEFb mediate, respectively, phosphorylation of Serine 5 and 2 (Ser5P, Ser2P), which control early transcription events at promoters. During transcript elongation, Ser5P levels decrease and Ser2P levels increase (Phatnani et al. 2006; Buratowski, 2009; Rahl al et al. 2010; Koga et al. 2015; Harlen et al. 2016). The RNAPII CTD is also target of PRMT5 (an arginine methyl-transferase that monomethylates or symmetrically dimethylates histone and nonhistone proteins), which symmetrically dimethylates the CTD arginine residue R1810 (Zhao et al. 2016). This R1810me2s modification recruits the survival motoneuron protein (SMN) to RNAPII elongation complexes. SMN, in turn, binds to a DNA-RNA helicase that resolves R-loops in TTS (transcription termination sites) and is important for proper termination and splicing of RNAPII transcripts (Zhao et al. 2016). Besides methylating RNAPII, PRMT5 acts on transcription through various histone methylations, which can activate or repress transcription according to the particular modification type. The repressor activity involves symmetrical dimethylation of R8 and R3 on histones H3 and H4, respectively, which leads to enhanced DNA methylation and chromatin compaction. Moreover, PRMT5 participates in transcriptional repressor complexes that include Histone Deacetylases (HDACs) and/or DNA methyl transferases (DMNTs) (Lacroix et al. 2008; Zhao et al. 2009; Karkhanis et al. 2011; Greenblatt et al. 2016; Blanc et al. 2017; Zhang et al. 2018; Antonysamy et al. 2017). PRMT5 functions instead as transcriptional activator by dimethylating R2 on histone H3 (Di Lorenzo et al. 2011; Liu et al. 2020). Thus, it is not surprising that it has multiple roles in cell biology - such as the regulation of neural differentiation, Golgi trafficking, and stemness maintenance - and is deregulated in a variety of tumors (Zhou et al. 2010; Chittka et al. 2012; Stopa et al. 2015; Scaglione et al. 2018; Banasavadi-Siddegowda et al. 2018). PRMT5 and PRMT5 inhibitors have gained a strong interest for developing new treatments for cancer (Friesen et al. 2002; Bedford et al. 2009, Beketova et al. 2021). We reported that MYC associates with PRMT5, enhancing PRMT5-dependent symmetrical dimethylation of H4 on R3 (H4R3me2s), and that PRMT5, in turn, regulates MYC activity (Mongiardi et al. 2015; Favia et al. 2019). The finding that PRMT5 has a key role in transcription termination suggested to us that MYC as well might be involved in transcription termination, through PRMT5. In the present work, we employ MYC inhibition by shRNA and by Omomyc to investigate MYC role in regulating RNAPII post-translational modifications, RNAPII distribution, and gene expression in cancer cells. Our study is suggestive of another control level exerted by MYC, which makes increasingly evident its role of master controller of gene expression, involved in RNAPII activation, elongation, termination, and pre-mRNA splicing.

## Results and discussion

### MYC induces RNAP II symmetrical dimethylation of R1810

To gain insights into a potential MYC role in transcription termination, we investigated by gain and loss of function experiments its ability to influence symmetrical dimethylation of RNAPII R1810, a key modification that is catalyzed by PRMT5 and regulates termination. To this end we employed Omomyc, a MYC specific shRNA, and the PRMT5 activity inhibitor EPZ015666. We transfected HEK293T recipient cells with a FlagMYC expression construct, together or not with MYC shRNA and FlagOmomyc expression plasmids. Following or not immunoprecipitation with RNAPII antibody, the protein extracts were analyzed by Western blotting with antibodies specific for PRMT5, SMN, R1810me2s (symmetrical dimethylation of RNAPII R1810), H4R3me2s (**Fig. 1 A,B**). We found that ectopic MYC expression strongly enhanced RNAPII R1810 symmetrical dimethylation, which was almost totally abolished by co-transfection with MYC-specific shRNA, as shown by the immunoprecipitations and the related densitometry histogram in **Fig. 1A,B**. Co-transfection of the Omomyc expression plasmid - which, by itself, affected R1810 dimethylation only weakly - strongly blunted the R1810 dimethylation increase caused by MYC ectopic expression. To verify that PRMT5 was required for the R1810me2s increase caused by over expressed MYC, HEK293T cells were transfected with the FlagMYC vector and treated or not with the PRMT5 inhibitor EPZ015666 24 hours after transfection. The PRMT5 inhibitor prevented the MYC dependent increase of R1810 dimethylation (**Fig. 1C**).

**Figure 1.**
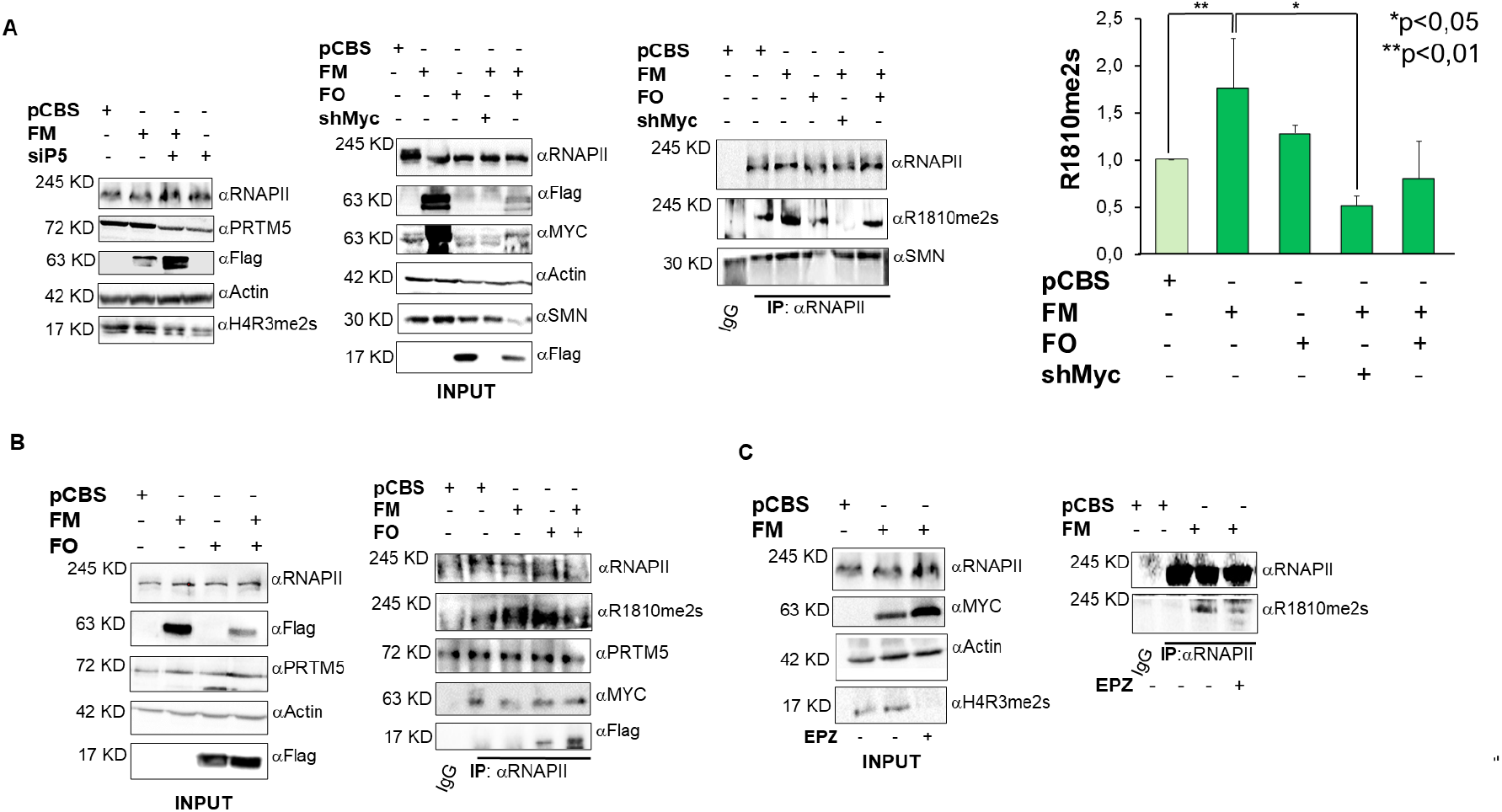
MYC ectopic expression enhances RNAPII R1810 symmetrical dimethylation via PRMT5. **A)** Left western blot: HEK293T cells transfected with a pool of siRNAs against PRMT5 and the day after with pCBS-FlagMYC plasmid. Immunoprecipitations were performed by a RNAPII antibody. The enhancement of H4R3me2s by MYC is sensitive to PRMT5 expression level. **A)** Middle and right blots, and **B)**: HEK293T cells transfected with pCBSFlagMYC, pCBSFlagOmomyc and a shRNA against MYC, alone or in combination. After 48 h, immunoprecipitations were performed by a RNAPII antibody. PRMT5, SMN, MYC and Omomyc co-precipitated with RNAPII. While ectopic MYC expression strongly increased RNAPII R1810 dimethylation, MYC shRNA and Omomyc inhibited such an increase (blots and densitometry histogram). Omomyc alone weakly induced R1810me2s. **C)** HEK293T: cells transfected with pCBSFlagMYC. The day after, cells were treated for 24 h with 5 μM EPZ01566 PRMT5 inhibitor or control vehicle, thereafter immunoprecipitation was performed. EPZ015666 impaired symmetrical dimethylation of H4R3 and the MYC-dependent increase of R1810me2s. Each bar in the histogram represents mean ± SEM. ***p-value 0,001, *p-value 0,05 repeated measures oneway ANOVA. Abbreviations. FM: FlagMYC; FO: FlagOmomyc; EPZ: EPZ015666; siP5: siPRMT5.

The immunoprecipitations in **Fig. 1A,B** indicate that PRMT5, SMN, MYC, and Omomyc are associated with RNAPII. In this regard, MYC associates with several proteins that regulate RNAPII activity (Gomez-Roman et al. 2003; Arabi et al. 2005; Rahl et al. 2010; Kaur, Cole et al., 2013; Campbell, White et al. 2014; WB et al. 2015; De Pretis et al. 2017). MYC and Omomyc functionally interact with PRMT5 (Mongiardi et al. 2015), which regulates transcription termination via SMN recruitment. All this confirms that MYC may participate in RNAPII complexes that regulate different aspects of transcription, termination included.

### MYC inhibition decreases RNAPII symmetrical dimethylation in cancer cells

MYC over-expression is a common feature of cancer cells. To evaluate MYC action on RNAPII symmetrical dimethylation, we took advantage of two cancer cell types with high MYC levels: the glioblastoma stem cell (GSC) line named Brain Tumor 168 (BT168) and the Burkitt lymphoma line RAMOS, in which the *myc* gene is under control of the immunoglobulin heavy chain promoter, (De Bacco et al. 2012; Dalla Favera et al. 1982; Bemark et al. 2000). The two cell lines were stably transduced with a doxycycline-inducible, FlagOmomyc (FO) expressing lentivirus (Galardi et al. 2016). As shown in **Fig. 2A,C** doxycycline treatment of BT168FO cells and RamosFO cells led to a strong reduction of RNAPII R1810me2s. The same result was obtained upon doxycycline treatment of BT168 cells stably transduced with a lentivirus expressing a doxycycline inducible shRNA against MYC (**Fig. 2B**). In both cell types the decrease of RNAPII symmetrical dimethylation was paralleled by a decrease of SMN binding to RNAPII, as expected.

**Figure 2.**
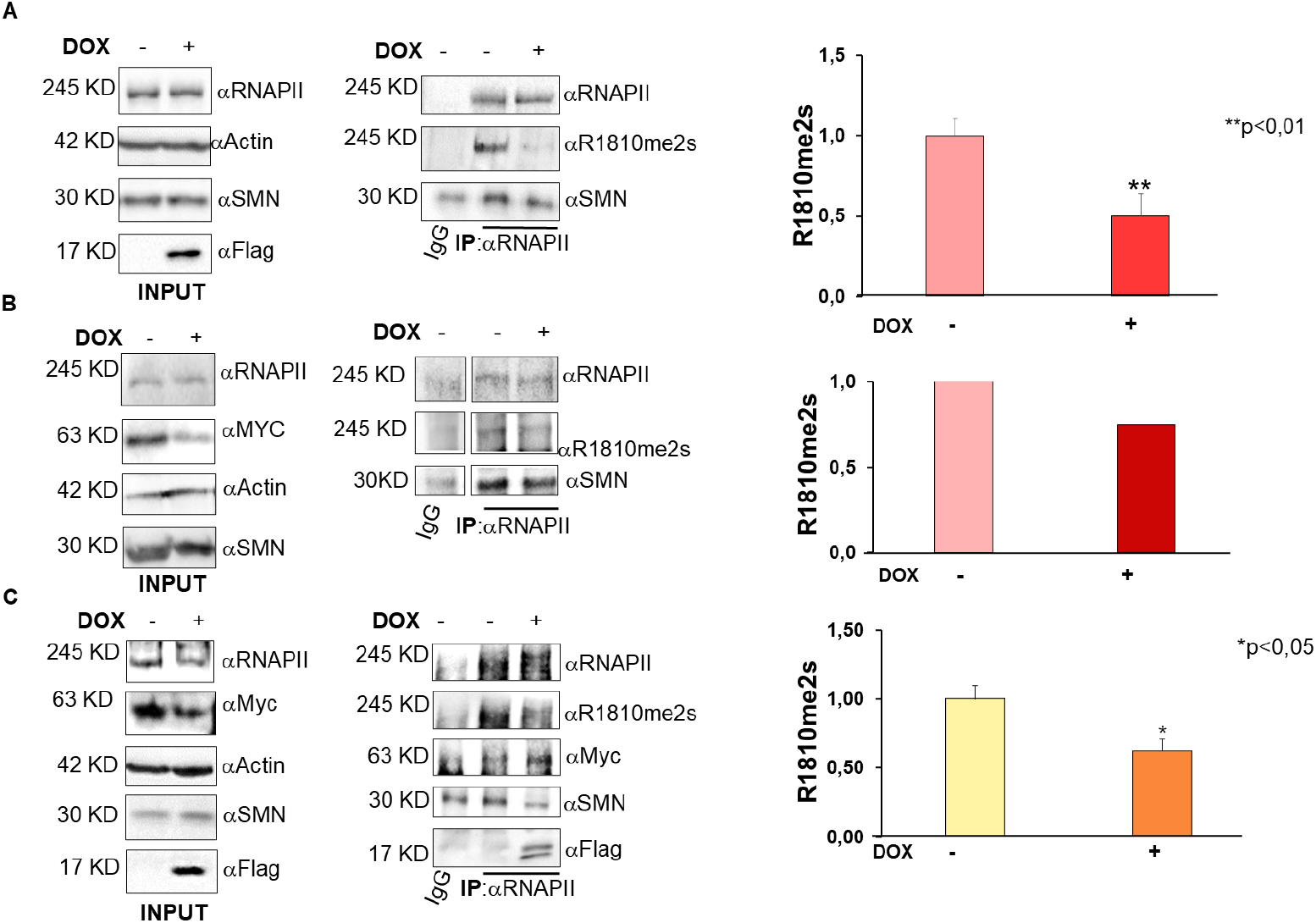
*MYC inhibition* by Omomyc and a shRNA *affects RNAPII R1810 symmetrical dimethylation in cancer cells*. Cells were treated with doxycycline for 24 hrs. Thereafter, immunoprecipitation was performed by means of a RNAPII antibody, followed by immunoblotting with RNAPII symmetrical dimethylation specific, and other antibodies. **A)** BT168FO cells. R1810 symmetrical dimethylation was severely impaired in the presence of FlagOmomyc. SMN binding to RNAPII also decreased. **B)** BT168 cells infected with a lentivirus encoding a doxycycline inducible shRNA against MYC. The shRNA strongly decreased RNAPII symmetrical dimethylation and SMN recruitment. **C)** RamosFO cells. Also in this cancer cell line, MYC inhibition led to a decrease of RNAPII symmetrical dimethylation and SMN recruitment. Densitometry histograms are shown to the right of each panel. Each bar represents mean ± SEM. ***p-value 0,001*p-value 0,05 paired t-tests.

Our data show that MYC sustains RNAPII symmetrical dimethylation and SMN recruitment, and that these actions are efficiently restrained by Omomyc and by a MYC shRNA. Overall, the findings that MYC interacts with PRMT5, strongly increases RNAPII R1810 symmetrical dimethylation, and affects SMN recruitment in different cell types, clearly suggest a novel role of MYC in regulating termination of transcripts by RNAPII.

### MYC inhibition decreases RNAPII Serine 2 phosphorylation and modulates MYC and MAX expression in cells expressing high MYC levels

The transition between initiation and productive elongation is elicited by Ser5 phosphorylation in RNAPII CTD, followed by the elongation-specific Ser2 phosphorylation. MYC binds to p-TEFB and regulates transcriptional pause release. Therefore, we asked whether Omomyc might affect Ser2 phosphorylation, thus influencing transcription elongation rate and mRNA expression. First, HEK293T cells were transfected with FlagMYC, FlagOmomyc, and pSLIK-shMYC plasmids, either alone or in combination. Immunoblotting analyses with RNAPII Ser2P specific antibodies showed that MYC ectopic expression strongly enhanced Ser2 phosphorylation (**Fig. 3A**), as expected. In parallel, MYC overexpression caused an increase of the p-TEFB component CDK9 (**Fig. 3A**).

**Figure 3.**
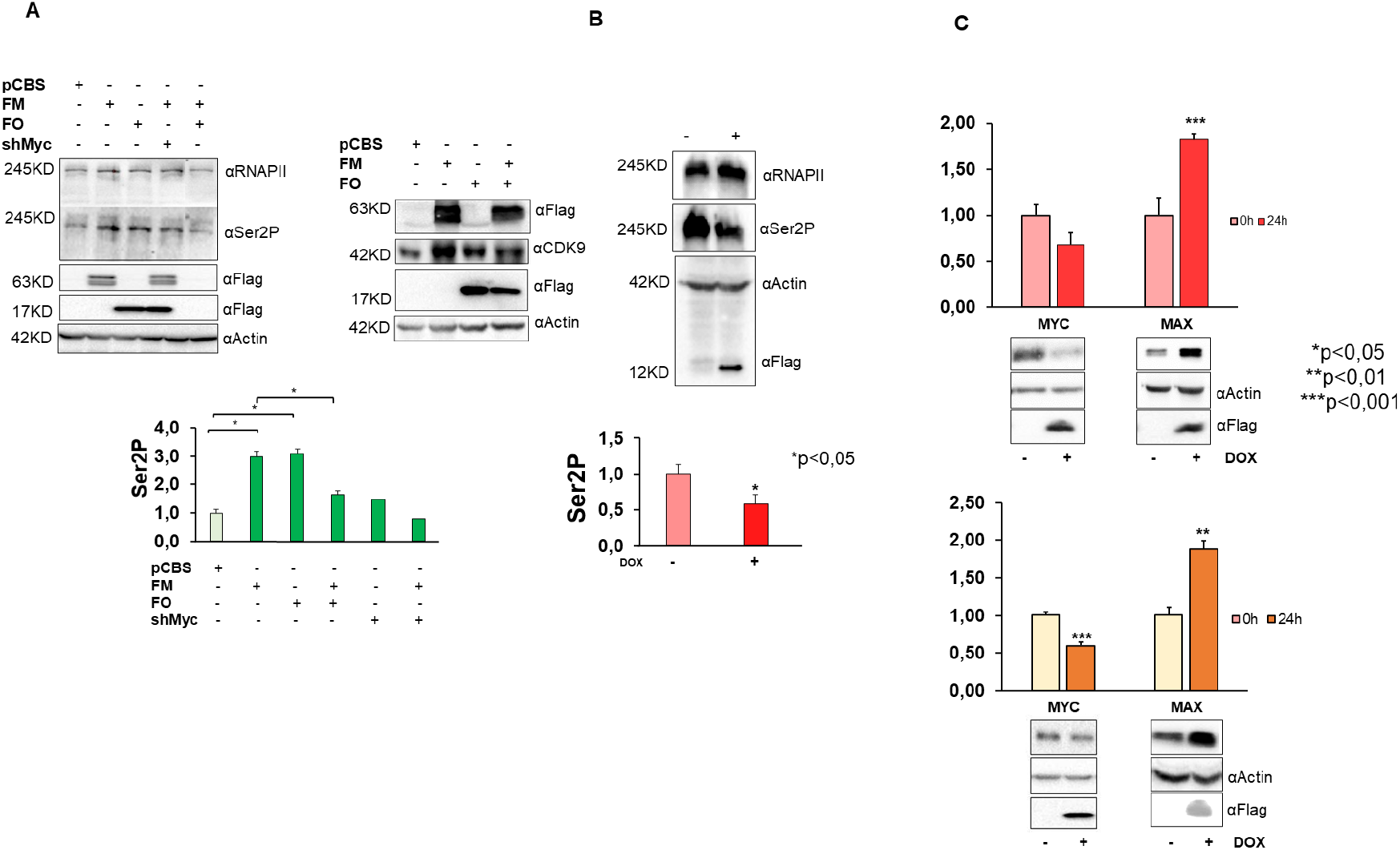
MYC inhibition impairs RNAPII Ser2 phosphorylation and increases Max protein. Representative immunoblots and densitometry histograms for quantifications of RNAPII Ser2P, MYC, and MAX expression in HEK293T, BT168FO, and RamosFO cells. **A)** HEK293T cells transfected with pCBSFlagMYC, pCBSFlagOmomyc, and MYC shRNA plasmids - alone or in combination - and probed with RNAPII Ser2P antibodies. **B)** BT168FO cells treated or not for 24 hrs with doxycycline, and probed with RNAPII Ser2P antibodies. **C)** BT168FO (top) and RamosFO cells (down) treated or not with doxycycline for 24 h, and probed with MAX and MYC antibodies. p-value** 0,01; p-value*** 0,0009 (mean ± SEM). Statistical analysis performed by paired t-test.

Omomyc expression as well enhanced Ser2 phosphorylation. In co-transfection experiments, instead, Omomyc caused a significant reduction of the levels of RNAPII Ser2P phosphorylation observed in the presence of over-expressed MYC (**Fig. 3A**, left panel). An even stronger Ser2P reduction was obtained by cotransfection with MYC shRNA (**Fig. 3A**). In BT168FO cells, RNAPII Ser2P decreased upon Omomyc induction (**Fig. 3B**). Altogether, data indicate that Omomyc may act on the transition between initiation and elongation by affecting Ser2P levels.

MYC overexpression in cancer cells contributes to the formation of a high number of MYC/MAX dimers that invade transcriptionally active chromatin sites (Nie et al. 2012; Lin et al. 2012; Guo et al. 2014; Sabò et al. 2014; Wolf et al. 2015). This causes as well a reduction of MAX homodimers, which attenuate the binding of MYC to specific (E-boxes) and non specific DNA sequences (Gu et al. 1993; Lindeman et al. 1995; Maltais et al. 2017). Omomyc affects such mechanisms by binding to chromatin as homodimers that largely displace MYC from DNA and can compromise DNA binding of MAX/MAX complexes as well (Savino et al. 2011; Jung et al. 2017; Galardi et al. 2016). We asked whether Omomyc might affect the expression of MYC and MAX as well. We found that its induction in BT168 and Ramos cells decreased the expression of MYC protein, in parallel with an increase of MAX (**Fig. 3C**). In HEK293T cells - which have low MYC levels compared to cancer cells like Ramos and BT168 - Omomyc did not significantly influence MYC expression (**Fig. 1A**, middle).

In conclusion, Omomyc appears to function by a multiplicity of ways: modulating MYC and MAX DNA binding and protein levels, inducing changes of RNAPII methylation, and perturbing the MYC interactome (Savino et al. 2011; Lourenco et al. 2021).

### Omomyc specifically represses direct MYC target genes

To further examine how Omomyc may influence the glioblastoma stem cell transcriptome, we measured by RNA-seq the mRNA output changes consequent to 24 and 48 h Omomyc induction in BT168FO cells. Genes that are MYC target and strongly expressed (FPKM <=10 in at least one condition) are shown in **Fig. 4 A,B.** As expected, the number of differentially expressed genes was higher in cells treated longer (**Fig. S1**). A 48 h treatment led to 2228 differentially expressed genes, 1606 of which were upregulated and 622 were downregulated (**Fig. 4 A,B, Fig. S1** and **Table S1**).

**Figure 4.**
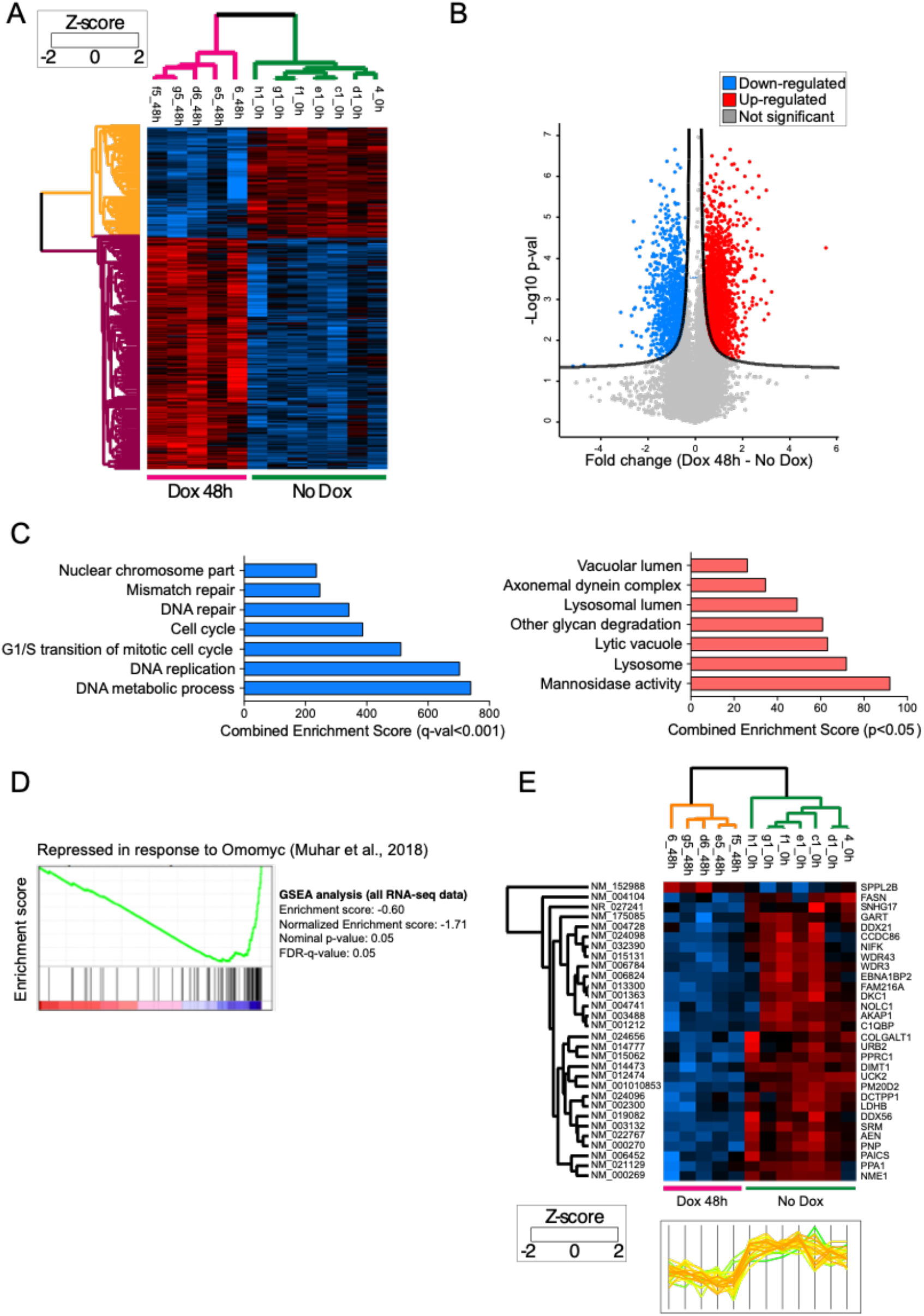
RNA sequencing and pathway analysis. **A,B)** RNA sequencing analysis shows 2228 genes that are differentially expressed in BT168FO cells upon 48 h Omomyc induction: 1606 up-regulated and 622 down-regulated. **C)** Functional enrichment analysis of GO terms in down-regulated (left) and up-regulated (right) genes upon 48 h Omomyc induction. The barplot show the top enriched GO biological processes, cellular component and molecular functions. **D)** GSEA analysis with enrichment score that confirmed the down-regulation of the Muhar signature of direct MYC targets genes upon Omomyc induction. **E)** Heatmap with profile plot of a subset of differentially expressed genes that are part of the Muhar signature. The subset includes genes strongly expressed - FPKM at least 10 - and differentially expressed, at p-value threshold < 0.05. All of them, except one, were downregulated upon Omomyc induction.

To elucidate the meaning of differentially expressed gene sets, we performed functional enrichment analyses by means of GO, KEGG, Reactome, WikiPathways. While we found little significance at 24 h DOX treatment - probably due to the lower number of differentially expressed genes (**Fig S1** and **Table S1**) - various pathways were significantly enriched after 48 h DOX treatment, coherently in the different analyses (**Fig. 4C**). Downregulated genes were enriched for gene ontology (GO) terms related to DNA metabolic process, DNA replication, DNA repair and cell cycle, in agreement with the view that MYC primarily acts as a transcriptional activator controlling metabolic and biosynthetic processes (Muhar et al. 2018; Kress et al. 2015). These GO terms were also correlated with MYC expression in a variety of cancer cell lines (Dang et al. 1999 and 2012; Lin et al. 2012; Fiorentino et al. 2016; Galardi et al. 2016; Lorenzin et al. 2016; Muhar et al. 2018). The pathways enriched among the upregulated genes were related to cell and amino acid metabolism, and lysosome activity. Altogether, these analyses indicate that Omomyc specifically influences the expression of genes and pathways that are regulated by MYC.

The differential expression observed following Omomyc induction is expected to be largely the consequence of MYC inhibition. Nevertheless, our gene expression analyses - done after relatively long perturbations of 24 and 48 h - cannot discriminate between direct and indirect effects. Furthermore, there is no consensus on direct regulatory functions of MYC, as several authors argue that MYC is a transcriptional activator and repressor of selected gene subsets, whereas others have suggested that MYC acts as a general transcriptional amplifier (Lin et al. 2012; Nie et al. 2012; Sabò et al. 2014; Walz et al. 2014; Lorenzin et al. 2016). To investigate Omomyc effect on the expression of genes directly regulated by MYC, we resorted to the signature of 100 direct MYC targets - conserved in a variety of cancer cell lines - described by Muhar and coworkers (Muhar et al. 2018). Such *Muhar signature* genes present a high level of MYC binding to the promoter, and their expression level in a given cell line correlates well with MYC amount in that cell line. We thus investigated whether Omomyc influenced the expression of this annotated gene set. We found that as many as 91 out of 100 *Muhar signature* genes were expressed in BT168FO cells (at FPKM>1), almost all significantly downregulated by Omomyc, as shown by the negative score in the Gene Set Enrichment Analysis (GSEA) and by the heatmap and profile score of the signature genes (**Fig 4 D,E**). This corroborates the hypothesis that Omomyc specifically represses the expression of authentic, direct MYC target genes.

### Omomyc affects RNAPII density at promoter and termination sites

MYC binds to most active promoters enhancing transcription pause release and elongation (Huang et al. 2021; Rahl et al. 2014). It also increases RNAPII R1810 symmetric dimethylation (Fig. 1), which regulates transcription termination by RNAPII (Zhao et al. 2016). Omomyc forms dimers that compete with MYC for DNA binding - causing a 50-60% reduction of MYC binding to promoters in BT168 cells (Galardi et al. 2016) - and restrains RNAPII symmetrical dimethylation, involved in transcription termination. This suggested to us that Omomyc might affect RNAPII density at promoters and/or terminator regions and influence gene expression at least partly in this way. In order to assess the influence of Omomyc on RNAPII distribution at promoter and terminator regions, we analyzed RNAPII ChIP-seq data of BT168FO cells treated or not with Dox, focusing on the genes that displayed promoter binding by MYC and were highly expressed (FPKM>=10). RNAPII density on either promoter and terminator regions of DOX treated versus control cells presented a clear linear correlation (**Fig 5 A,B**). We found that Omomyc induction led to a strong increase (1.5 - 2 fold) of the amount of RNAPII bound to promoter and termination sites (**Fig. 5 A,B**). The increased density at promoters may be explained by the consideration that Omomyc restrains transcriptional pause release and may thus cause an accumulation of RNAPII at promoters. The increased density at terminators may be explained by the impaired R1810 symmetrical dimethylation, which affects termination and may lead to RNAPII accumulation at termination regions of active genes (Zhao et al. 2016). RNAPII relative density - expressed as the ratio between terminator and promoter density - was instead insensitive to Omomyc, remaining unchanged or showing only a very slight decrease in DOX treated versus untreated cells (**Fig. 5C**).

**Figure 5.**
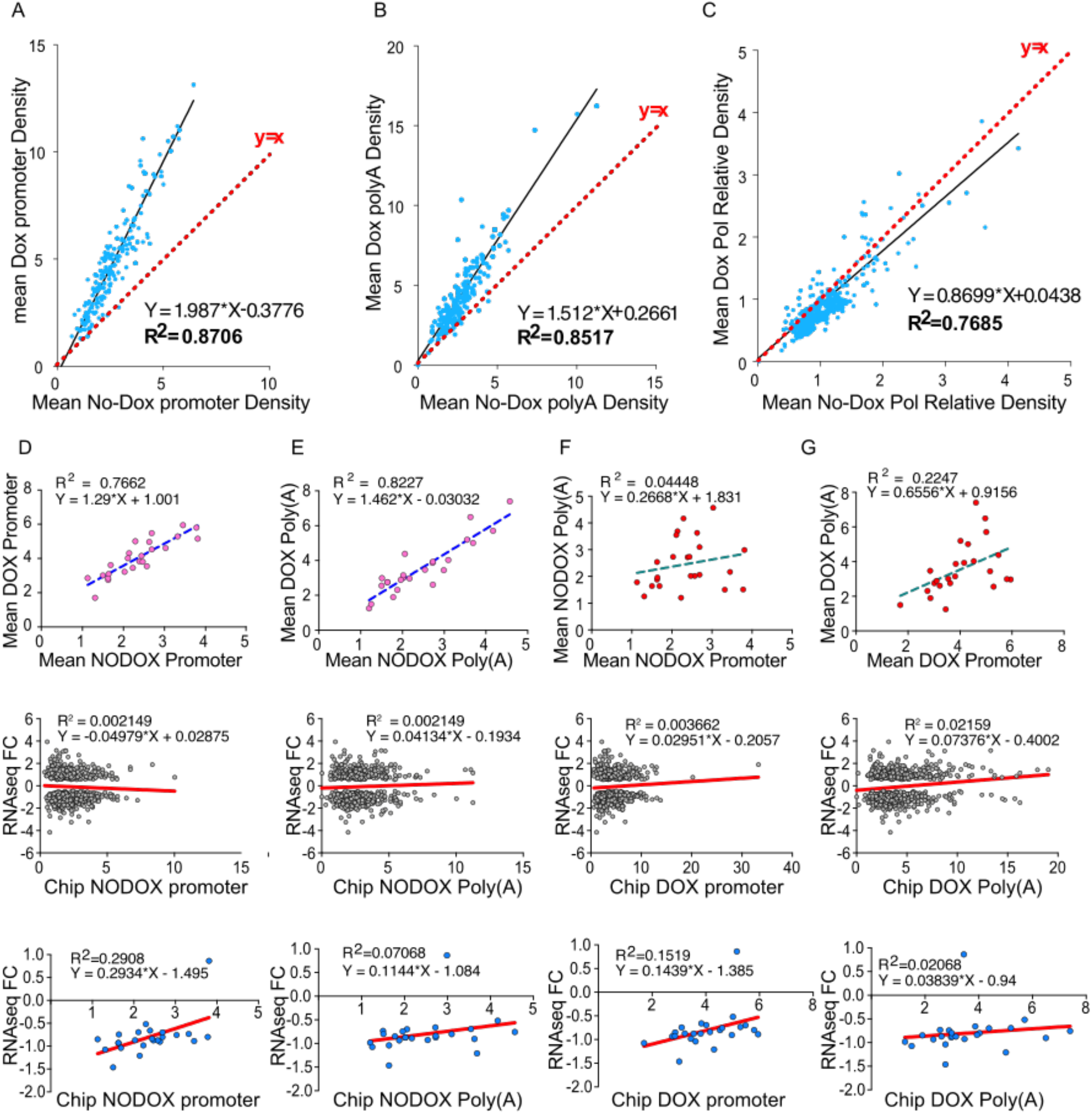
MYC inhibition affects RNAPII density at promoter and terminator regions. Correlation scatter plots of RNAPII occupancy and mRNA fold change (FC) in genes strongly expressed (FPKM>=10) in at least one condition and with a significant Fold Change (p-value threshold <0.05) in BT168FO cells treated or not with DOX for 48 h. **First two rows (A-G):** RNAPII density at promoter and PolyA regions; **A-C:** all MYC target genes; **D-G:** Muhar signature genes (the same as in Fig. 4, panel E). The scatter plots show an increased RNAPII occupancy upon Omomyc induction at both promoter and termination sites, slightly higher at promoters. **Third and fourth rows:** comparison between RNAseq Fold Change and CHIPseq density at promoters and terminators. **Third row:** all MYC targets; **fourth row:** Muhar signature genes (as in **Fig. 4**,).

**Figure 6.**
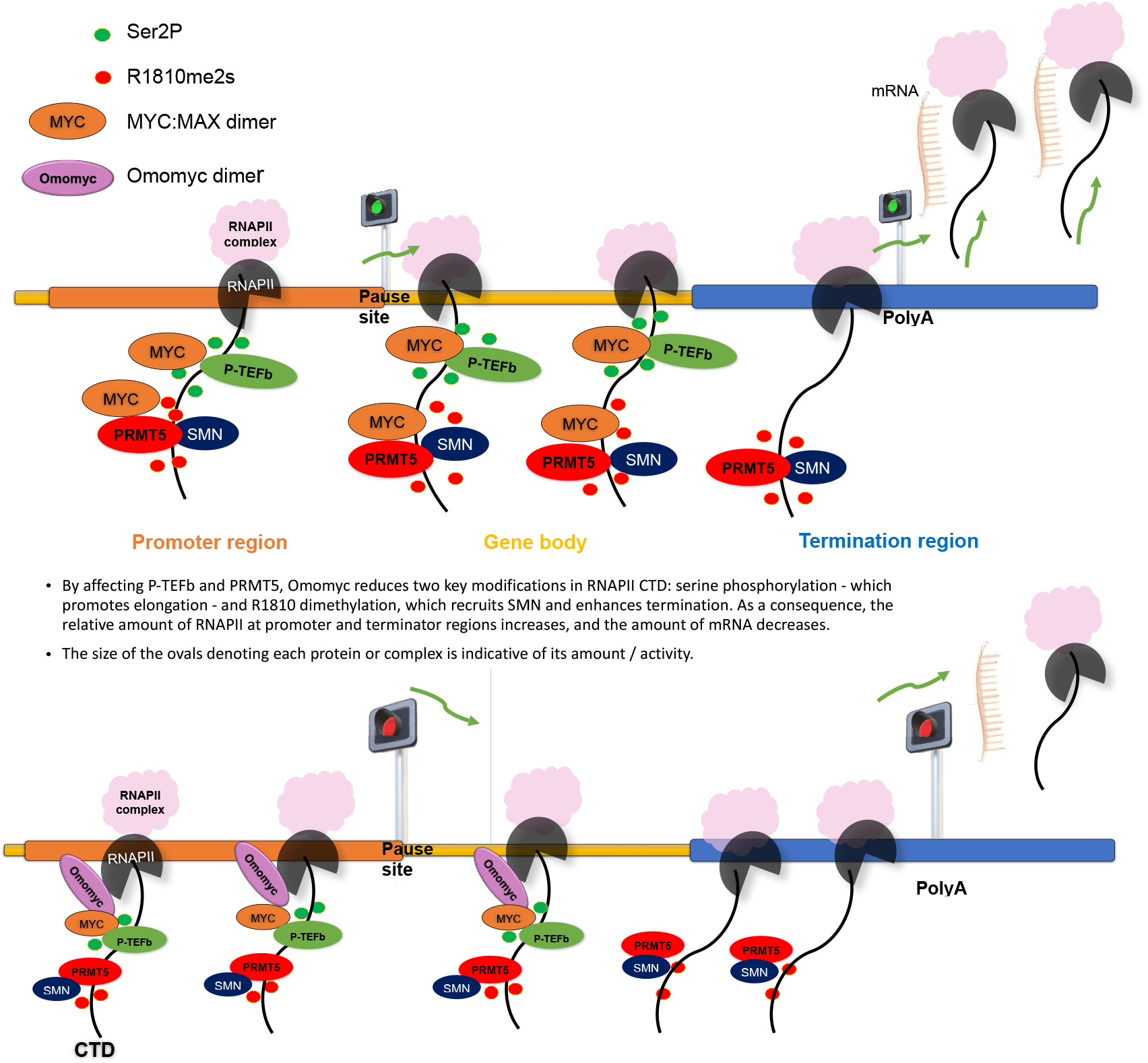
Model.

This trend was also present in the *Muhar signature* of MYC target genes (**Fig 5 D-G**), with higher values of RNAPII density at promoters and terminators in DOX treated cells. As to mRNA expression levels, *Muhar signature* genes were significantly down-regulation by Omomyc induction. The correlation analysis for all MYC target genes (**Fig. 5, third row**) did not show a strong link between mRNA expression changes and RNAPlII occupancy. So, while Omomyc induction leads - either directly and indirectly - to global expression changes of MYC target genes in BT168FO cells, such changes cannot be solely explained by Omomyc-induced changes in RNAPII density at terminator and/or promoter sites, but would involve modulation by additional factors, the action of which might be directly influenced by Omomyc.

In summary, Omomyc impairs MYC binding to promoters, perturbs the MYC interactome (see also Savino et al. 2011), and restrains MYC dependent enhancement of transcriptional pause release and R1810 dimethylation. All this contributes to an adjustment of RNAPII distribution and to renormalization of the expression of genes that are deregulated as a consequence of MYC overexpression - such as those governing the GSC phenotype (Galardi et al. 2016) - by molecular mechanisms that are yet to be clarified.

## Author contributions

FS performed most experiments, contributed to analyze genomic data and to writing the manuscript; AP performed bioinformatics data analyses and contributed to writing; AF did some of the biochemistry experiments; CS designed the MYC shRNA and supervised the experiments with Ramos cells; DZ and JG provided the R1810me2s antibody and gave some suggestions; BI codesigned and supervised many of the biochemistry experiments, and contributed to writing; SN designed and supervised the whole work and wrote the manuscript.

## Acknowledgments

We thank Dorothy Zhao and Jack Greenblatt (Donnelly Centre, University of Toronto) for donating the RNAPII R1810me2s antibodies and for suggestions

## MATERIALS AND METHODS

### Cell lines and culture

BT168 Glioblastoma stem cells (GSC) cells were previously described by De Bacco et al. 2012. Cells were grown as neurospheres in serum-free medium, DMEM/F-12 (SIGMA, St.Louis, Mo, USA) supplemented with B-27^™^ Supplement (50X), 1% penicillin/streptomycin, 2mM Glutamine (Thermo Fisher Scientific), 10 ng/mL EGF and bFGF (Life Technologies, Carlsband, CA). Burkitt’s lymphoma Ramos cells were characterized by Dalla Favera et al. 1982. Cells were cultured in RPMI-1640 medium supplemented with 10% FBS (Thermo Fisher Scientific), 1% penicillin/ streptomycin, 2mM Glutamine (Thermo Fisher Scientific). HEK293T cells were cultured in Dulbecco’s Modified Eagle Medium (DMEM, SIGMA, St. Louis, Mo, USA), supplemented with 10% FBS (Thermo Fisher Scientific), 2 mM Glutamine and penicillin/streptomycin (Thermo Fisher Scientific). Cells harboring a doxycycline inducible Flag-Omomyc were obtained by lentiviral infection. BT168FO and Ramos FO cells were treated respectively with 0.25 μg/mL and 0.1 μg/mL doxycycline (SIGMA) to induce Omomyc. BT168shMYC1# cells were obtained by transduction with an inducible lentivirus expressing a short hairpin RNA for MYC; they were treated with 0.25 μg/mL doxycycline to induce shMYC. HEK293T cells were treated with 5uM EPZ01566 (1:1000) PRMT5 inhibitor. Cells were harvested 48 h after treatment and the inhibition of PRMT5 activity was tested with Immunoblots for H4R3me2s.

### Lentiviral infection

The lentiviral plasmid pSLIK-FO was already described (Galardi et al. 2016). The lentiviral plasmid pSLIK-shMYC1# (sh sequence TGCTGTTGACAGTGAGCGAAAGATGAGGAAGAAATCGATGTAGTGAAGCCACAGATGTACATCGATTTCTCCTCATCTTCTGCCTACTGCCTCGGA) was engineered by cutting pSLIK-FO using PacI and SnaBI to cut away Gateway platform. The fragment PacI-SnaBI was purified. PCR from GEPIR (all-in-one shRNA-vector; Fellmann et al. 2013) for TRE3G-EGFP-mir30E band inserting the SnaBI and PacI sites. The fragment TRE3G-EGFP-mir30E was purified and cloned in pSLIK-PacI-SnaBI vector. pSLIK-SnaBI-mir30E-PacI was cutted with SnaBI for re-inserting RRE and Flag sequence. The final vector pSLIK-shMYC co-express hygromycin resistance gene and Tet-transactivator rtTA3. Lentiviruses were prepared by co-transfecting (Lipofectamine 2000 reagent, Thermo Fisher Scientific) HEK293T cells with pSLIK-Flag-Omomyc and packaging plasmids PLP1, PLP2 and pMD VSV-G diluted in Opti-MEM (Thermo Fisher Scientific). The medium was removed after 12-24h and replaced with 4mL of fresh growth media. Supernatants were collected every 24 h between 48 to 72 h after transfection, pulled together and concentrated by ultracentrifugation in a Beckman SW-28 rotor for 2h at 25000 rpm, 4°C. For infection, the 2-5 × 105 cells were seeded in 35mm dishes and infected the following day in the presence of 4 μg/mL polybrene. BT168FO cells were selected with 50–200 μg/mL hygromycinn B (Sigma), Ramos FO cells with 400-800 μg/mL. After selection, Flag-Omomyc and shMYC expression was assessed by western blots.

### Transfection

Flag-Omomyc (pCbsFlag-Omomyc), Flag-MYC (pCbsFlag-MYC), pSLIK-shMYC plasmids were transfected using Lipofectamine 2000 (Thermo Fisher Scientific) according to the manufacturer’s instructions. 25nM PRMT5-siRNA or control siRNA (GE Dharmacon, SiRNA-SMART pool) were transfected with PepMuteTM siRNA transfection reagent (SignaGen Laboratories) according to the manufacturer’s instructions. Cells were harvested for 48h after transfection.

### Immunoprecipitation

Immunoprecipitation was performed with RIPA buffer (140mM NaCl, 10mM Tris pH 7.6-8.0, 1% Triton, 0.1% sodium deoxycoholate, 1mM EDTA, containing protease inhibitors (Roche) and benzonase (SIGMA) (Zhao et al. 2016). 10 - 20 × 106 cells were lysed on ice for 25 minutes by vortexing and forcing them through a 27-gauge needle, at least 10 times. After centrifuging at 13000 rpm for 15 min at 4°C, the supernatant was incubated with 10μL-25μL of protein A/G beads (Thermo Fisher) and 1-2μg of antibodies for 4h to overnight. The samples were washed 3 times with RIPA buffer and boiled in SDS gel sample buffer. To detect R1810me2s modification on RNA polymerase II (RNAPII), alkaline phosphatase (Roche) treatment (5μL) at 37°C for 30 min was performed for RNAPII immunoprecipitated samples before boiling (Zhao et al. 2016).

### Immunoblotting

Proteins were resolved in 6-8-10 or 12% polyacrilammide gels and transferred to PVDF (Bio-Rad, Hercules, CA,USA) or nitrocellulose membranes (GE Heath Care, Little Chafont, Buckinghamshire, UK) for 2h at 250 mA on ice or over-night at 30V. Filters were blocked in phosphate buffered saline plus 0.1% Tween-20 (PBST, SIGMA) added with 10% non-fat dry milk, for 1 hour and half at room temperature (RT). Primary antibodies were incubated over-night (O/N) at 4 °C, according to the concentration recommended by the manufacturer, in PBST plus 2.5%-5% non-fat dry milk. After three 10 minutes washes, filters were incubated for 1 hour at RT with either goat-anti rabbit (1:5000) or goat-anti mouse (1:2000) horseradish peroxidase (HRP)-conjugated secondary antibodies (Merck Millipore, Darmstadt, Germany). Blots were developed using SuperSignal West Pico Chemiluminescent Substrate or Femto Maximum Sensitivity (Thermo Fisher Scientific). Images were captured with a Chemidoc XRS+ (Bio-Rad, Hercules, CA, USA) and quantified using ImageJ software. Anti-MYC (9E10 and N-262), anti-CDK9, anti-RNAPII (8WG16) antibodies were from Santa Cruz Biotechnologies, anti-H4R3me2s, anti-PRMT5 antibodies were from Abcam; anti-Flag antibody was from SIGMA. Anti-R1810me2s was courtesy of J. F. Greenblatt’s lab – University of Toronto (Zhao et al. 2016). Anti-dimethyl-Arginine Antibody, symmetric (SYM10) was from Merk. Anti-RNA polymerase II CTD repeat YSPTSPS (phospho S2) was from Abcam. Anti-β-Actin-peroxidase was from SIGMA.

### Chromatin Immunoprecipitation (ChIP), ChIP-seq and RNA-seq

Samples for ChIP and ChIP-seq assays were prepared and analyzed according to Myers Lab ChIP-seq Protocol v041610 (http://myers.hudsonalpha.org/documents/) and MAGnify Chromatin Immunoprecipitation System protocol (Invitrogen). Antibodies used: MYC (sc-764Z, Santa Cruz Biotechnologies), MAX (c-197X, Santa Cruz), RNAPII (sc-899X, Santa Cruz), RNAPII phospho Ser5 (ab5131, Abcam), RNA Pol II phospho Ser2 (ab24758, Abcam and 3E19, Active Motif), Flag (F1804, Sigma). For RNA-seq, 2μg total RNA purified by PureLinkRNA Mini Kit (Life Technologies) was used. ChIP-seq and RNA-seq libraries were prepared at Istituto di Genomica Applicata (IGA; www.appliedgenomics.org/) according to Illumina TruSeq DNA and TruSeq RNA Sample Preparation Guides. Samples were sequenced through Illumina HiSeq 2000 e 2500.

### Data processing and bioinformatics analysis

Data were processed as described in Galardi et al. 2016. For ChIP-seq analysis, 50-bp reads were mapped to hg19 human reference genome (UCSC Genome Browser) using Bowtie (Langmead et al. 2009) version 0.12.7 allowing three mismatches; reads with multiple best matches were discarded. Peak calling was through MACS (Zhang et al. 2008) 1.4.2 with 10-4 P-value cut-off. The RefSeq transcript annotation of hg19 was used for computing intersections between peaks and promoters. Binding enrichment to promoters was calculated by the normalized number of ChIP-seq reads as Reads Per Million (RPM). In case of multiple TSSs, those with the highest enrichment were chosen. Motif enrichment analysis was performed by Giulio Pavesi (University of Milan) through Pscan-ChIP (Zambelli et al. 2013). Seqminer v. 1.3.3 was used to calculate distribution around TSSs. The RAP RNA-Seq pipeline (https://bioinformatics.cineca.it/rap/) - including quality controls, adaptor trimming and masking of low-quality sequences, tophat2, bowtie, and CuffLinks 2.2 - was used to reconstruct the transcriptome (hg19 reference) and calculate expression values as FPKM (Fragment per Kilobase Million per genes). Data have been analyzed using DESeq2 R package (Love et al. 2014), considering genes with a FPKM > 0. Differentially expressed genes between treated (24 hours and 48 hours) and untreated samples with an adjusted p-value<0.05 were taken as up-regulated (log2 fold change > 0) or down-regulated (log2 fold change < 0). Up- and down-regulated genes were separately used for Gene Ontology Enrichment Analysis using EnrichR (Kuleshov et al. 2016) and Gene Set Enrichment Analysis using GSEA (Subramanian et al. 2005). Data and figures were further analyzed using in-house R scripts and Perseus tool (Tyanova et al. 2016). Comparisons between MYC and Omomyc occupancy and gene expression (FPKM), were performed calculating the average values for groups of 100 genes (bins) and correlated by a scatter diagram. The linear regression model was used to assess the correlation between transcript levels in NODOX versus DOX cells. RNAPII distribution, at TTS versus TSS regions, was evaluated using ChIP-seq data. Density reads, counted as RPKM, for each gene, at promoter (1500 nt) and termination (4200 nt) regions was calculated dividing the number of reads by the total number of reads obtained from each sequencing per condition (-DOX and +DOX), and by the length of the features. Data were normalized by their INPUT. Gene Set Enrichment Analysis (GSEA, http://www.broad.mit.edu/gsea/index.html) was used to determine whether an a priori defined set of genes shows statistically significance, according to the differences between -DOX and +DOX experimental conditions (phenotypes). In details, the RNA-Seq dataset files – consisting of experiments in triplicate for each time point of DOX treatment – containing two labeled phenotypes (-DOX and +DOX) were prepared in TXT format: -DOX included all 0h time point (1° phenotype), while +DOX included from 4h to 48h of DOX treatment (2° phenotype). The expression dataset was compared with several genes sets either exported from GSEA-MsigDB database or homemade. The gene sets contained the gene set name and the list of included genes. A gene set file was in GMX or GMT format. GSEA software calculated an enrichment score (ES) describing the degree to which a gene set was overrepresented at the extremes (top or bottom) of the entire ranked list of data set - where genes are ranked according to the expression difference between -DOX and +DOX conditions. The Enrichment Score ES was calculated by walking down the list. The value statistically increased when it found genes present in the gene set and decreased when genes were not present. The magnitude of ES was dependent on the correlation of each gene with the phenotype. The proportion of false positives was evaluated by calculating False Discovery Rate FDR-q value. Refseq IDs were mapped onto gene symbols using biormaRt R tool (Durinck et al. 2009). Analyses have been performed on the mean value of promoter and termination sites for each condition, calculated as the mean value of two independent replicates normalized per million of mapped reads. Correlation analysis between RNA sequencing and Chip sequencing has been performed using the log2 fold change value for RNA-seq and the mean promoter/termination site for Chip-seq. RNAPII density has been calculated as the ratio between the mean termination site and the mean promoter site values.

### Statistical analysis

Statistical analyses were performed by using the GraphPad Prism version 5.0d (GraphPad, La Jolla, CA) and Excel (Microsoft Excel, version 2018). All histograms represent the mean ±SEM of data obtained in 3 or more independent experiments. Statistical significance was determined by one-way repeated-measures ANOVA or paired t-test. The box plot p-values were calculated by paired Wilcoxon signed-rank tests. Regression lines were estimated using linear regression models. For genomic data, differential expression was assessed by CuffDiff2, as well as by Fold-Change thresholds, and Gene Set Enrichment Analysis (GSEA: www.broadinstitute.org/gsea/) subdividing MYC targets and non-MYC targets in groups of 500 genes.

### Data availability

ChIP-seq and RNA-seq data used in this study are accessible via Gene Expression Omnibus (GEO http://www.ncbi.nlm.nih.gov/geo/), with accession identifier GSE86519.

