## Supplemental Figure 1 for "A new insight into MYC action: control of RNA polymerase II methylation and transcription termination"

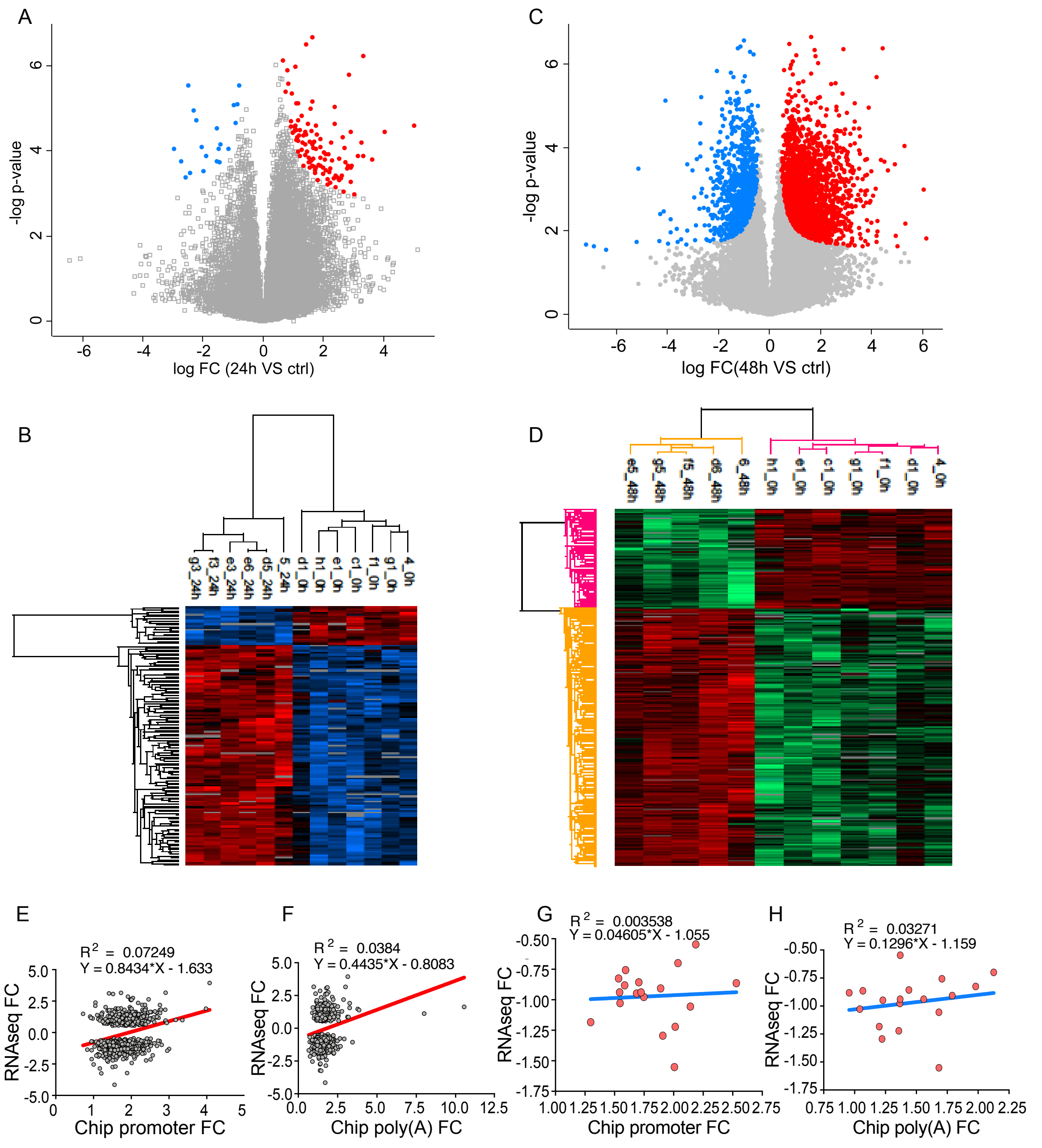


***Figure S1. RNA-seq analysis of BT168FO glioblastoma cells***
***A-D*** *mRNA expression at 24 h vs 48 h DOX treatment:* ***A,B****: 24 h DOX,* ***C,D****: 48 h DOX;* ***A,C*** *volcano plots,* ***B,D*** *heatmaps.* ***E-H*** *mRNA expression FC vs RNAPII occupancy FC at promoter and PolyA regions:* ***E,F****: all MYC targets;* ***G,H****: Muhar signature genes (direct MYC targets).*
